# Evaluating performance and applications of sample-wise cell deconvolution methods on human brain transcriptomic data

**DOI:** 10.1101/2023.03.13.532468

**Authors:** Rujia Dai, Tianyao Chu, Ming Zhang, Xuan Wang, Alexandre Jourdon, Feinan Wu, Jessica Mariani, Flora M. Vaccarino, Donghoon Lee, John F. Fullard, Gabriel E. Hoffman, Panos Roussos, Yue Wang, Xusheng Wang, Dalila Pinto, Sidney H. Wang, Chunling Zhang, PsychENCODE consortium, Chao Chen, Chunyu Liu

## Abstract

Sample-wise deconvolution methods have been developed to estimate cell-type proportions and gene expressions in bulk-tissue samples. However, the performance of these methods and their biological applications has not been evaluated, particularly on human brain transcriptomic data. Here, nine deconvolution methods were evaluated with sample-matched data from bulk-tissue RNAseq, single-cell/nuclei (sc/sn) RNAseq, and immunohistochemistry. A total of 1,130,767 nuclei/cells from 149 adult postmortem brains and 72 organoid samples were used. The results showed the best performance of dtangle for estimating cell proportions and bMIND for estimating sample-wise cell-type gene expression. For eight brain cell types, 25,273 cell-type eQTLs were identified with deconvoluted expressions (decon-eQTLs). The results showed that decon-eQTLs explained more schizophrenia GWAS heritability than bulk-tissue or single-cell eQTLs alone. Differential gene expression associated with multiple phenotypes were also examined using the deconvoluted data. Our findings, which were replicated in bulk-tissue RNAseq and sc/snRNAseq data, provided new insights into the biological applications of deconvoluted data.

## Introduction

Brain transcriptome is essential for studying brain biology and related disorders, but important cell type information can be obscured when bulk tissue is used for the data production. Several brain projects have generated valuable transcriptomic resources from human brains, such as GTEx(*1*), PsychENCODE(*2*), CommonMind(*3*), Brainspan(*4*), and ROSMAP(*5*). However, most of the existing transcriptomes are from bulk tissue, which are mixtures of many different cell types, and gene regulatory mechanisms are known to vary across brain cell types, obscuring the cellular mechanisms underlying bulk tissue expression changes.

Sorted-cell RNAseq and single-cell/nuclei (sc/sn) RNAseq(*6, 7*) offer solutions for profiling brain transcriptome at the cell-type resolution but with several limitations. Cell sorting relies on marker genes, which are not always available. The specificity of such marker genes is frequently a concern. A combination of several marker genes only can sort limited cell types. The data from sc/snRNAseq is sparse due to the limited RNA input from each cell(*8*). Sc/snRNAseq data suffer from the large number of zero values, which are called “dropout events”. Moreover, it is challenging to discriminate between two possible causes of dropouts: biologically true zero expression and technical random missing data(*9*). The presence of dropouts may result in potential problems in gene expression quantification. Another limitation is the high cost of sc/snRNAseq. Even though multiplexing methods have been developed to simultaneously profile cells from numerous samples(*10*), using sc/snRNAseq in large-scale studies typically requiring hundreds of subjects, such as disease association and expression quantitative trait loci (eQTL) mapping(*11*), can be cost-prohibitive.

Computational algorithms for cell deconvolution have been developed to estimate cell proportions. These algorithms can be classified into two types: supervised deconvolution uses prior information from a cell-type reference data to facilitate the estimation of the cell proportions of each cell types in bulk-tissue samples, while unsupervised does not need a reference. This study focused on evaluating supervised deconvolution methods.

The performance of methods for estimating cell proportions has been previously evaluated (*12-16*). Studies have evaluated the accuracy of estimated cell proportions with data from the brain and other tissues. Methods like DSA(*17*), OLS, CIBERSORT(*18*), dtangle(*19*), and MuSiC(*20*) showed good performance in these evaluations. Additionally, the effect of cell type marker gene selection, covariates, data transformation and normalization, and cell subtypes on cell deconvolution has been evaluated, which provided guidelines for data processing before cell deconvolution.

Estimated cell proportions have been used for cell-type studies but with limitations. Cell proportions were used to represent cell types for case-control comparison(*21, 22*). However, proportional changes in cell types are just one aspect of possible changes. Disease-related changes also involve cell-type-specific gene expressions(*23-26*). Cell proportions have also been used to map eQTLs associated with cell types, which are called cell-type interaction eQTLs(*27*) (ieQTLs). The genetic regulators that were associated with gene expression when the cell proportion varied were mapped. ieQTL has two major limitations. Firstly, ieQTLs are not necessarily specific to cell types. They may refer to other cell types with positive or negative correlation with the cell proportion of the target cell type. Secondly, the power of ieQTL mapping is low. Less than 50 ieQTLs were reported for neurons with 15,201 samples (*27*), which is much less than what standard eQTL can map on bulk tissues. Therefore, there is still the need to discover expression changes associated with diseases and eQTLs from cell-type expression data.

Cell-type gene expressions can be deconvoluted from bulk-tissue expression data. Methods have been developed to estimate cell-type expressions for each sample, such as bMIND(*28*), swCAM(*29*), and TCA(*30*). We call this sample-wise deconvolution of gene expression. These methods use expression references from sc/snRNAseq or sorted-cell expressions. For example, bMIND used the Bayesian model and Markov Chain Monte Carlo to estimate expression for each gene in the cell types of each sample. The cell-type expressions of the individual samples will enable eQTL mapping and differential expression analysis in cell types. The deconvoluted data can cover the majority of genes in bulk tissue and is less sparse than sc/snRNAseq data. It makes the large sample study of cell-type expression affordable since bulk tissue data is either ready to use or can be generated at a relatively low cost.

The methods for estimating sample-wise cell-type expressions have been partially evaluated with major blind spots. The performance of bMIND, TCA, and swCAM has been evaluated in their original methodology papers. However, these studies used artificially-constructed pseudo-bulk data other than bulk-tissue data to benchmark their performance. Pseudo-bulks were constructed by simulating cell proportions and multiplying these proportions with expressions from sc/snRNAseq or sorted-cell expression data. Therefore, pseudo-bulk data is less complex than data from real bulk tissue(*31*). The differences among cell types in the pseudo-bulk are easier to be captured than those in bulk-tissue data. The benchmark conclusion based on the pseudo-bulk data may not apply to data from brain tissues. Head-to-head comparisons of all these methods on brain data have not been conducted to date. The downstream applications based on deconvoluted data, such as eQTL mapping and differential expression, have also not been evaluated to showcase the validity of deconvolution.

The current study aimed at evaluating the performance of algorithms for sample-wise deconvoluting cell proportions and cell-type expressions, as well as research applications based on the deconvoluted data. Specifically, we evaluated six commonly-used deconvolution methods for estimating cell proportions and three deconvolution methods for estimating the cell-type expressions of individual samples. Data from bulk-tissue RNAseq, sc/snRNAseq, and immunohistochemistry (IHC) of matched adult postmortem brains and brain organoids were used for evaluation. Downstream analyses of the deconvoluted results were also conducted, including their use in eQTL mapping, schizophrenia (SCZ) GWAS heritability enrichment, differential expression for Alzheimer’s disease (AD), SCZ, and brain development in cell types. Based on the evaluation, we recommended the best practice for brain transcriptome deconvolution.

## Results

### Benchmarking of sample-wise deconvolution methods with brain transcriptome data

To evaluate commonly-used deconvolution methods, we selected six methods (DSA, OLS, CIBERSORT, dtangle, MuSiC, and Bisque(*32*)) for estimating cell proportions and three methods (bMIND, swCAM, and TCA) for estimating cell-type expressions (Fig. 1). Bulk-tissue RNAseq, snRNAseq, and IHC data from ROSMAP were used as primary data for evaluation(*33*). Data from adult brains in CommonMind (CMC)(*34*) and brain organoids(*35*) were used for confirmation (Table 1). Cell proportions from IHC and sc/snRNAseq data were used as ground truth for evaluating the accuracy of estimated cell proportions. Gene expressions in sc/snRNAseq data were used as ground truth for evaluating the accuracy of estimated cell-type expressions. The root-mean-square error (RMSE) and Spearman correlation coefficient were used as evaluation metrics. After method evaluation, eQTL mapping, GWAS heritability enrichment, and differential expression analysis were performed on the cell-type expressions estimated by the best performing method. To further evaluate the quality of outputs of these deconvolution methods by actual applications, the eQTLs, explained GWAS heritability, and phenotype-associated genes derived from deconvoluted expressions were compared to corresponding results based on sc/snRNAseq and bulk-tissue data.

**Fig. 1.**
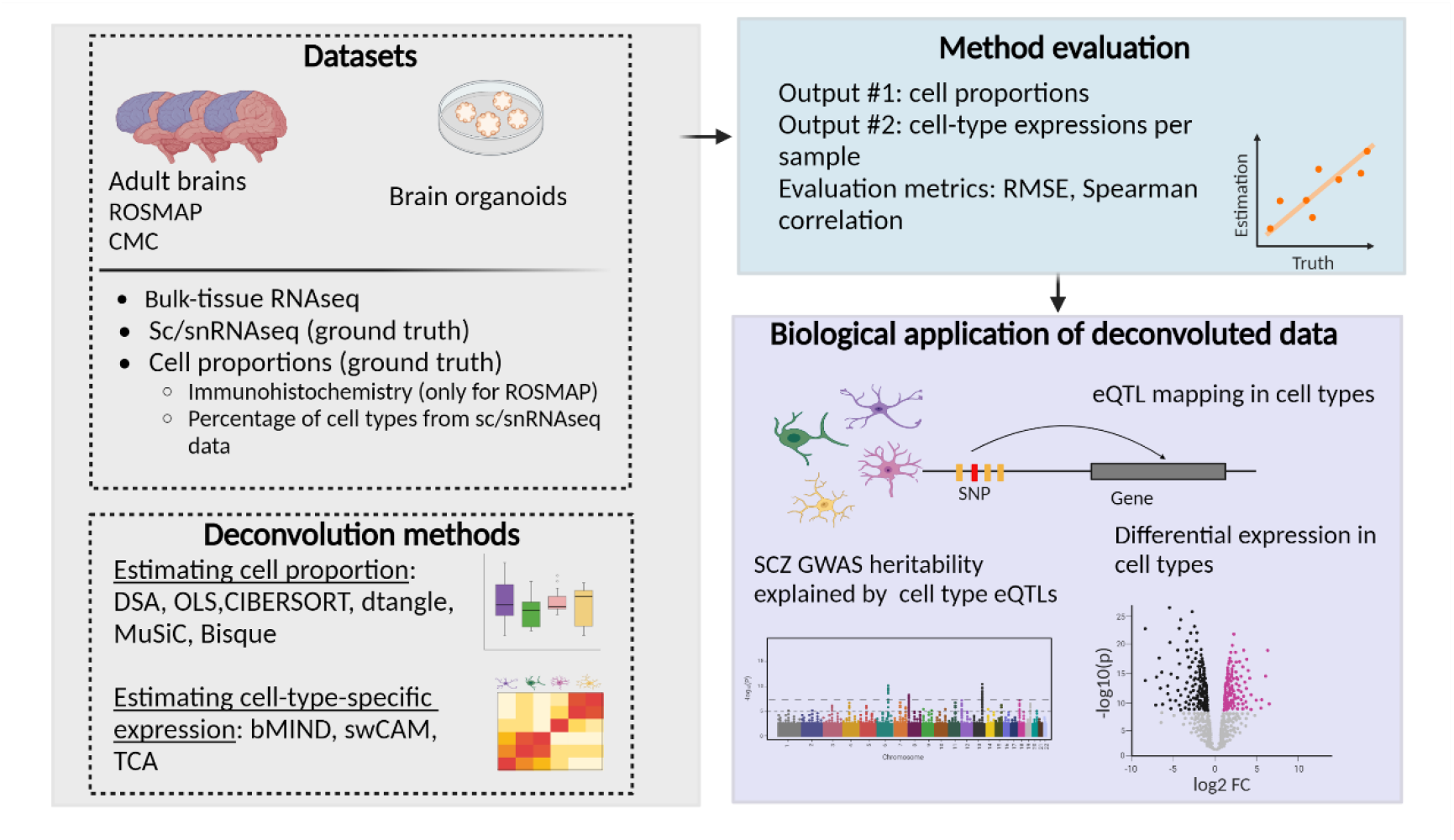
Study overview.

**Table 1.** Datasets used for evaluation

| Study | Brain region | Data type | Sample size | Number of cells | Number of cell types | Number of genes |
| --- | --- | --- | --- | --- | --- | --- |
| ROSMAP | PFC | Bulk-tissue RNAseq | 1,112 | - | - | 17,128 |
|  | PFC | snRNAseq | 48 | 69,611 | 8 | 17,926 |
|  | PFC | IHC | 49 | - | 5 | - |
| CMC | PFC | Bulk-tissue RNAseq | 572 | - | - | 25,774 |
|  | PFC | snRNAseq | 101 | 569,289 | 7 | 33,822 |
| Brain organoid | - | Bulk-tissue RNAseq | 130 | - | - | 20,125 |
|  | - | scRNAseq | 72 | 490,844 |  | 33,538 |

\*PFC: prefrontal cortex

### Evaluation of cell proportions estimated by deconvolution methods

The overall performance of six deconvolution methods (DSA, OLS, CIBERSORT, dtangle, MuSiC, and Bisque) for estimating cell proportions was evaluated with ground truth from matched samples. To ensure the deconvolution performance, the intersection of marker genes identified at the individual-cell level and the pseudo-bulk level was used to guide deconvolution (see details in methods). Using ROSMAP IHC data as ground truth (n of samples=49), dtangle, and OLS showed lower RMSE than other methods (Fig. 2A). dtangle also showed relatively low RMSE in CMC (n=94) and brain organoid (n=55) data. MuSiC and Bisque did not perform well, even though they are designed to use sc/snRNAseq data as a reference. Using cell proportions computed from sc/snRNAseq data as the ground truth, Bisque had the lowest RMSE in all three datasets. The accuracy of deconvoluted cell proportions in major cell types was better than that in minor cell types (Fig. 2B, Fig. S1). The RMSE increased sharply when the cell proportion was below 5%, such as in oligodendrocyte precursor cells (Opc), microglia, endothelial cells, and pericytes in adult brains. Similar results were observed using Spearman correlation as an evaluation metric (Fig. S2).

**Fig. 2.**
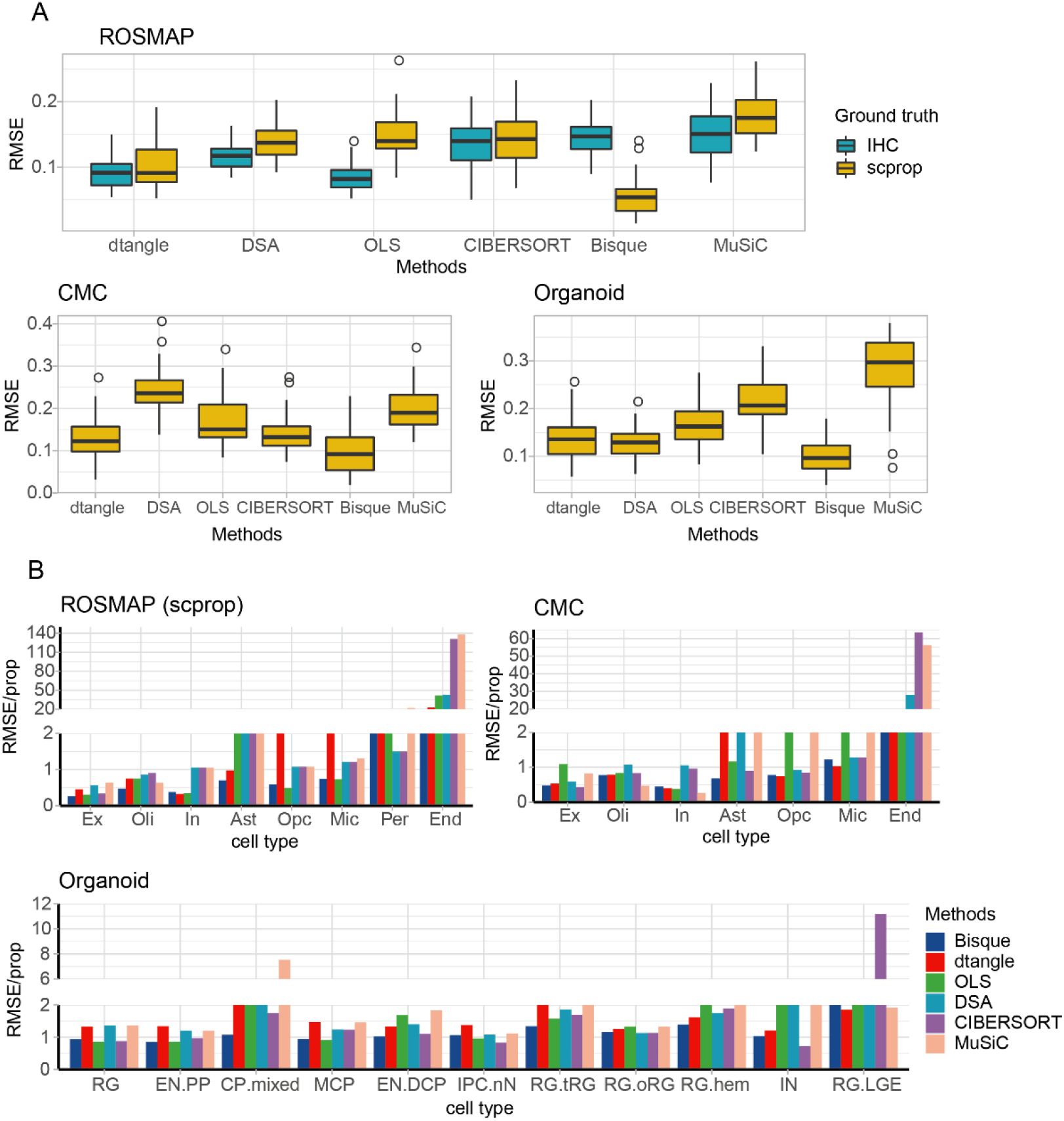
Assessment of cell proportions estimated by examined deconvolution methods. **(A)**. Sample-level RMSE values between estimated cell proportions and ground truth. IHC: immunohistochemistry; scprop: cell proportions calculated from sc/snRNAseq data, scprop = the number of cells of specific cell type/number of total cells. **(B).** Cell-type-level RMSE values between estimated cell proportions and ground truth data. RMSE values were normalized by the value of cell proportions to make them comparable across cell types. Cell types were ordered by cell proportions in a decreasing way. Ex: excitatory neurons, In: inhibitory neurons, Ast: astrocytes, Opc: oligodendrocyte precursor cells, Mic: microglia, Per: pericytes, End: endothelial cells; RG: radial glia, EN.PP: early born excitatory neurons of the pre-plate/subplate, CP.mixed: cortical plate mixed neurons, MCP: medial cortical plate, EN-DCP: dorsal cortical plate excitatory neurons, IPC-nN: intermediate progenitor cell or newborn neuron, RG.tRG: truncated radial glia, RG.oRG: outer radial glia, RG.hem: radial glia in cortical hem, IN: inhibitory neurons, RG-LGE: progenitors corresponding to a putative ventrolateral ganglionic eminence fate.

### Evaluation of sample-wise cell-type expressions estimated by deconvolution methods

The accuracy of sample-wise cell-type expressions deconvoluted by bMIND, swCAM, and TCA was evaluated using ground truth generated from sc/snRNAseq expressions of matched samples (n=35 for ROSMAP, n=94 for CMC, and n=55 for brain organoid). Cell proportions estimated by Bisque and dtangle were selected as input for the three methods, since they showed the best performance in the above evaluation for estimating cell proportions. bMIND showed the best performance for estimating cell-type expressions in all datasets, followed by swCAM (Fig. 3A). For bMIND, the averaged correlation coefficient between estimated expression and sc/snRNAseq data was 0.62 in ROSMAP data, 0.75 in CMC adult brain data, and 0.85 in brain organoid data. We did not observe a substantial difference in performances for estimating expressions of major and rare cell types (Fig. 3B). However, bMIND performed more steadily and overall better in major cell types than in minor cell types. The deconvoluted expressions by bMIND correlated with corresponding cell types in sample-matched sc/snRNAseq data, and they were less correlated with unrelated cell types (Fig. S3). A number of well-known marker genes were highly expressed in corresponding cell types, thus indicating that the deconvoluted data have good cell-type specificity (Fig. 3C).

**Fig. 3.**
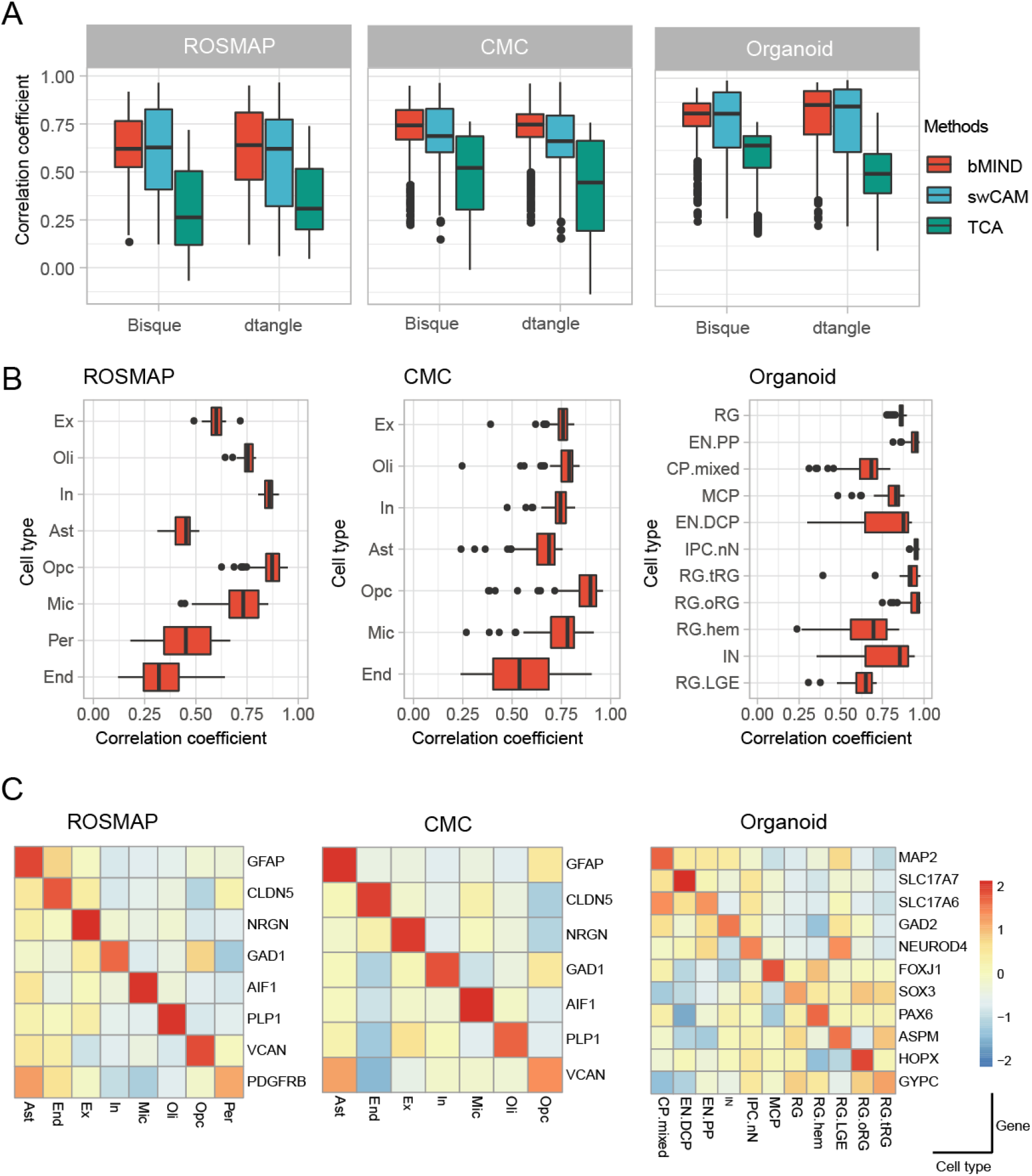
Assessment of sample-wise cell-type expressions deconvoluted from bulk-tissue data. **(A).** Overall assessment of methods for estimating cell-type expressions. Spearman correlations between deconvoluted data and sc/snRNAseq data from matched samples. The averaged expression by cell types was used as ground truth. **(B).** Cell-type-level assessment of methods for estimating cell-type expressions. Correlations between deconvoluted data by bMIND and sc/snRNAseq data were calculated for each cell type. Cell proportions estimated by dtangle were used for input. Cell types on the y-axis were ordered by cell proportions computed from sc/snRNAseq. **(C).** Assessment of cell type specificity in estimated expressions. The figure shows the expression of marker genes in deconvoluted data by bMIND.

### Cell-type eQTL mapping with deconvoluted sample-wise expression data

To identify SNPs that cis-regulate gene expression in specific cell types, cell-type eQTL mapping was performed for the association between genotypes and deconvoluted gene expression data of individual samples. The cell-type eQTLs identified with deconvoluted gene expression data were named deconvolution eQTLs (decon-eQTL). RNAseq data of 1,112 bulk-tissue samples of ROSMAP collection were deconvoluted. Cell proportions and cell-type expressions were estimated with dtangle and bMIND, respectively. Out of the 1,112 samples, 861 had genotype data and were used for decon-eQTL mapping. The effect of SNPs within a 1-megabase window around the transcription start site (TSS) of genes was tested. The numbers of input genes for decon-eQTL mapping ranged from 8,521 to 12,418 across all cell types. The number of input SNPs was 4,954,561 for all cell types. Effects of known and hidden covariates on deconvoluted expressions were corrected. A total of 1,088,634 to 2,245,945 decon-eQTLs were detected across eight cell types at a genome-wide significant level (FDR<0.05). To identify the independent effect in SNPs, a permutation test was performed for each gene. A total of 25,273 (4,541∼ 8,149) independent decon-eQTLs were identified at FDR<0.05 for eight cell types (Fig 4B). As expected, eQTL SNPs (eSNPs) were enriched around the TSS region of eQTL genes (eGenes) (Fig. S4). The numbers of detected decon-eQTLs were positively correlated with the proportions of cell types in the tissue (Fig. 4B). To test the robustness of identified decon-eQTLs, sample IDs were randomly shuffled before the eQTL mapping. The absence of significant eQTL in the shuffled data supported that the identified decon-eQTLs were not due to random noise (Fig. S5).

**Fig. 4.**
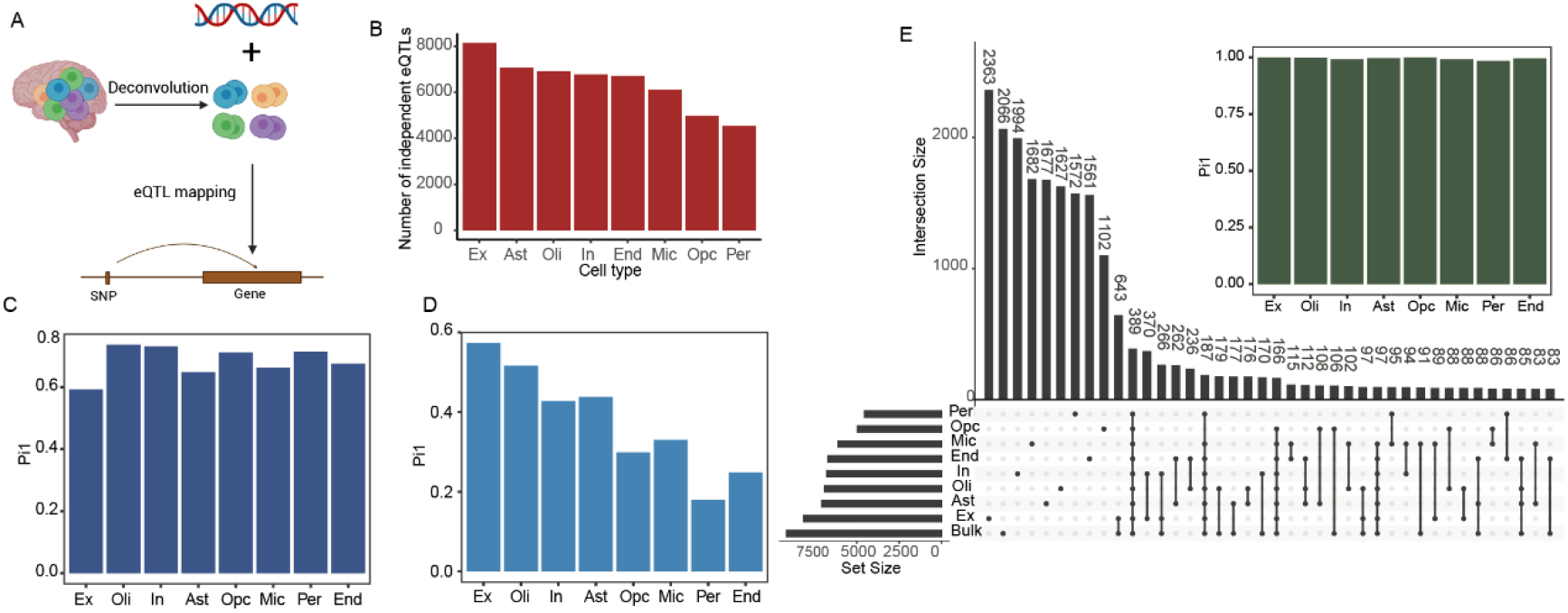
Cell-type eQTL mapping based on deconvoluted sample-wise expression data. **(A)** Illustration of decon-eQTL mapping. **(B)** The number of decon-eQTLs identified in different cell types at FDR<0.05 in the permutation test. **(C)** Pi1 statistics of decon-eQTLs in BrainGVEX decon-eQTLs and **(D)** eQTLs from snRNAseq study (Bryois et al.). **(E)** Comparison of decon-eQTLs and bulk-tissue eQTLs. The top barplot shows the Pi1 values of decon-eQTLs in bulk-tissue eQTLs. The bottom plot shows the intersections between decon-eQTLs and bulk-tissue eQTLs, as well as intersections of decon-eQTLs across various cell types.

Identified decon-eQTLs from ROSMAP data were replicated with another deconvoluted data from BrainGVEX(*36*). The same deconvolution and eQTL mapping procedures were performed on RNAseq data of 400 postmortem brain samples from BrainGVEX to obtain the decon-eQTLs. Across all eight cell types, 3,479 to 5,718 independent eQTLs were identified in the deconvoluted data from BrainGVEX at FDR<0.05. To measure the replication rate of ROSMAP decon-eQTLs in BrainGVEX data, Pi1 statistic(*37*), which is the proportion of true eQTL associations in the replication data, were calculated. The Pi1 of ROSMAP decon-eQTLs in BrainGVEX data was 0.59 ∼ 0.74 for the matched cell types (Fig. 4C). Decon-eQTLs of oligodendrocyte had relatively better replication than other cell types.

Cell-type eQTLs from snRNAseq data in Bryois et al.(*38*) were also used to replicate our decon-eQTLs. This replication study had performed genotyping and snRNAseq on 192 cortical samples. A total of 7,607 independent eQTLs across eight cell types were identified. Even though the replication data had less statistical power than our deconvoluted data, 17%∼57% of decon-eQTLs were replicated (Fig. 4D). eQTLs of excitatory neurons (Pi1=0.57) had higher Pi1 values than other cell types (averaged Pi1=0.38).

To illustrate the value of decon-eQTLs, we compared decon-eQTLs to bulk-tissue eQTLs from ROSMAP (Fig. 4E). Overall, decon-eQTLs had good replication in bulk-tissue data, with Pi1>0.95. The eQTLs that were significant at the cell-type level but insignificant at the bulk-tissue level were defined as cell-type-specific eQTLs. A total of 1,206 ∼ 3,006 (24.3% ∼ 36.89%) cell-type-specific eQTLs were identified in the deconvoluted data. Cell-type-specific eQTLs had Pi1 values of 0.17∼0.52 in single-cell eQTLs, which were similar to Pi1 values of decon-eQTLs that were shared with bulk-tissue eQTLs (Fig. S6). This demonstrated that a good proportion of eQTLs regulate gene expressions in a cell-type-specific way, and they can be detected by decon-eQTLs.

### Cell-type eQTLs enriched for the risk heritability in SCZ GWAS data

To test whether cell-type eQTLs are enriched for genetic risk heritability of SCZ, stratified linkage disequilibrium score regression (sLDSC)(*39*) was used to calculate the heritability of SCZ GWAS mediated by decon-eQTLs. Single-cell eQTLs and bulk-tissue eQTLs were also included for comparison. Decon-eQTLs explained more SCZ GWAS heritability (averaged h2 = 37%) than single-cell eQTLs (averaged h2 = 6%) for all cell types (Fig. 5A). Bulk-tissue eQTLs explained 49% of SCZ GWAS heritability. Integrating decon-eQTLs and bulk-tissue eQTLs increased the explained heritability to 63%, whereas the integration of single-cell eQTLs only resulted in an increase of heritability to 53%. The total proportion of explained heritability was correlated with the proportions of each cell type. To control the effect of SNP numbers, heritability was normalized by the number of decon-eQTLs, which was called enrichment. Decon-eQTLs of all cell types were enriched for SCZ GWAS heritability (Fig. 5B, P value<0.05). Decon-eQTLs of oligodendrocytes showed the strongest per-SNP enrichment across all cell types. The SCZ GWAS heritability was only significantly enriched in single-cell eQTLs from oligodendrocytes and excitatory neurons. Decon-eQTLs of most of the cell types showed higher enrichment of SCZ GWAS heritability than bulk-tissue eQTLs (Fig. 5B), indicating that some of the SCZ risk SNPs may affect gene expression in cell type-specific ways. Deconvolution analyses uncovered more such cell-type-specific regulations associated with the genetic risk of SCZ.

**Fig. 5.**
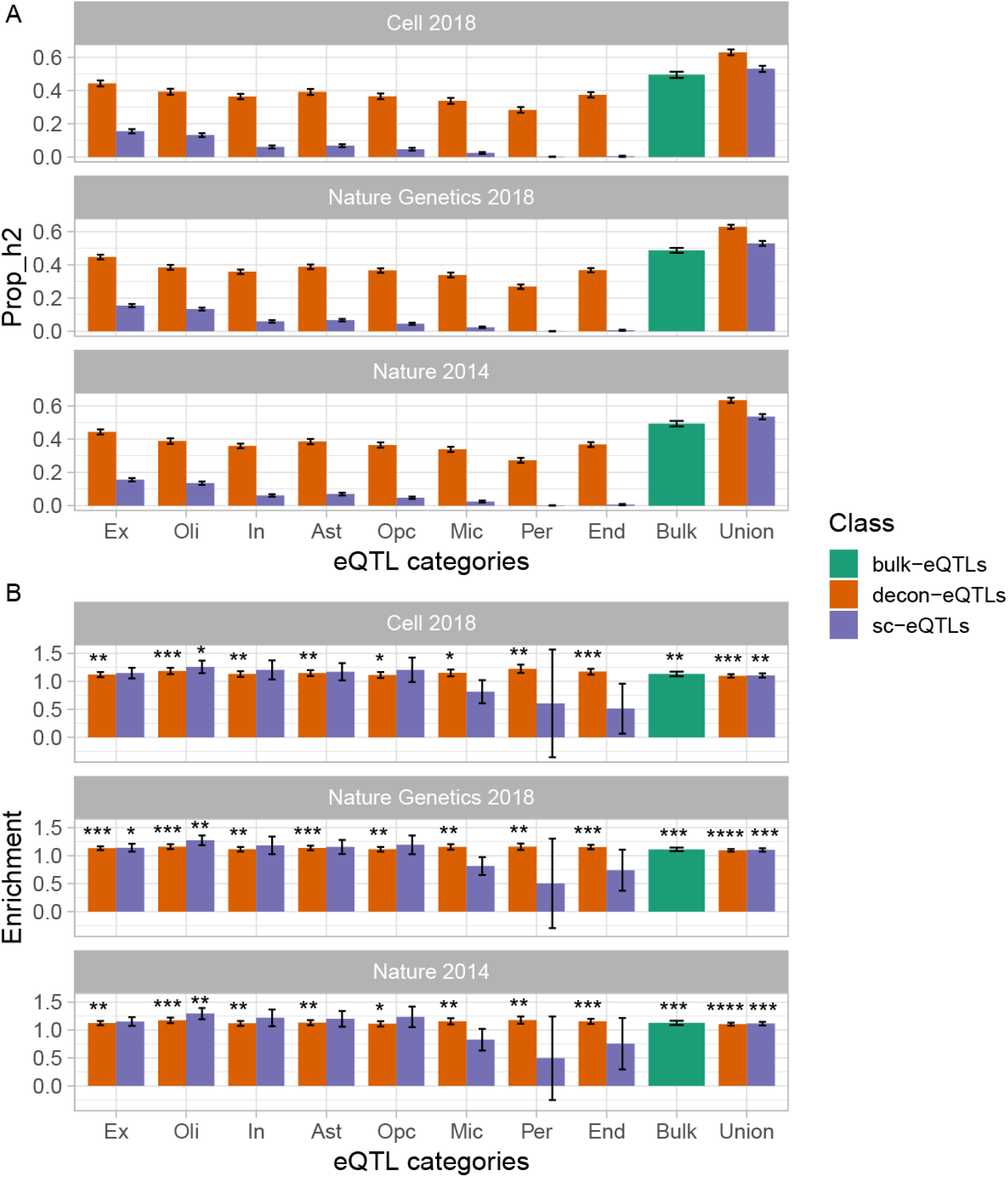
SCZ GWAS heritability explained by cell-type eQTLs and bulk-tissue eQTLs. **(A)** Total SCZ GWAS heritability (h2) explained by eQTLs. **(B)** SCZ GWAS heritability enrichment in eQTLs. Enrichment = h2/number of SNPs in each eQTL category.

### Identification of gene expression changes associated with disease and brain development within cell types

To identify genes associated with various phenotypes in specific cell types, differential gene expression analysis was conducted using the deconvoluted sample-wise expression data. Associations with AD, SCZ, and brain development modeled by organoids were tested in three deconvoluted datasets independently. More samples were included in AD (N_AD_=743, N_control_=367) and SCZ (N_SCZ_=246, N_control_=279) data. For AD and SCZ, the Wilcoxon signed-rank test was performed on the deconvoluted data. For brain development, the linear regression model was used to test the correlation between deconvoluted data and culture days of organoids (N_day0_=15, N_day30_=22, N_day60_=18). With a threshold of FDR<0.05, 4,419, 10,964, and 9,562 phenotypes-associated genes (PAGs) were identified for AD, SCZ, and brain development, respectively.

To test the reliability of PAGs identified from deconvoluted data, these PAGs were compared to those identified from bulk-tissue and sc/snRNAseq data (Fig. 6). In total, 81%, 49%, and 89% of PAGs for AD, SCZ and brain development, respectively, were replicated in bulk-tissue data. Among these PAGs, most of them (>95%) had the same direction of expression changes in bulk-tissue data. For AD and SCZ, less than 15% of PAGs overlapped with PAGs from snRNAseq data. However, 35% of development-related PAGs could be replicated in scRNAseq data. The possible explanation for the difference in replication rate in the three datasets was that the expression changes associated with brain development were larger than the changes associated with AD or SCZ (Fig. S7). The low replication rate with sc/snRNAseq data suggested that sc/snRNAseq data was underpowered to detect PAGs of small effect size.

**Fig. 6.**
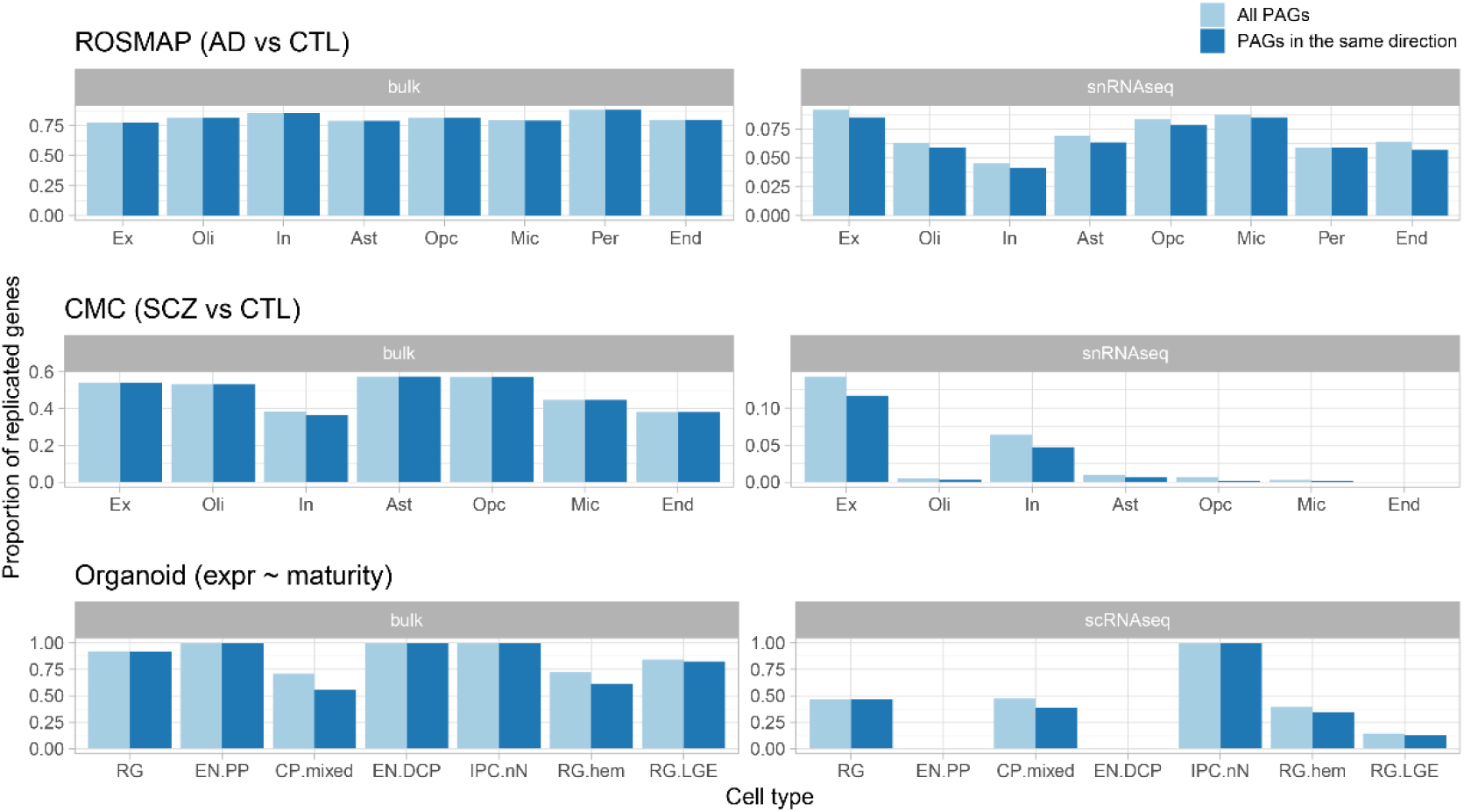
The proportion of phenotypes-associated genes (PAGs) replicated in bulk-tissue data (left panel) and sc/snRNAseq data (right panel). The light blue bar shows the proportions of replicated PAGs with FDR<0.05. The dark blue bar shows the proportions of replicated PAGs with FDR<0.05 and having the same direction of changes in replication data. expr denotes expression. The maturity of organoids was measured by the days of cell culture.

## Discussion

Using matched samples of bulk-tissue RNAseq, IHC, and sc/snRNAseq data, the performance of six methods for estimating cell proportions and three methods for estimating sample-wise cell-type gene expression was systematically evaluated. The transcriptome data used for evaluation were from adult brains and cultured brain organoids, providing data representative from different states of cell maturity and developmental processes. In addition, the results of eQTL mapping, SCZ GWAS heritability enrichment, and differential expression analysis based on deconvoluted data demonstrate the utility of deconvolution.

dtangle had better accuracy for estimating cell type proportions than other methods. Previous studies have benchmarked the performance of deconvolution methods for estimating cell type proportions with ground truth data created from simulated proportions. In those studies, dtangle showed good performance in Sutton et al(*15*) but poor performance in Avila Cobos et al.(*12*) One possible reason is the difference of pseudo-bulk simulation between studies. Sutton et al. constructed pseudo-bulks from 500 cells while Avila Cobos et al. used only 100 cells for each pseudo-bulk. Given that sc/snRNAseq data are remarkably sparse, 100 cells may not be representative of cell composition in bulk tissue. This inconsistency suggests the necessity of using real ground truth data in benchmarking studies. By using bulk-tissue RNAseq and IHC data from matched samples, dtangle was found to be the best deconvolution method for estimating cell proportions in this study. The excellent performance of dtangle was preserved in two replication data.

Deconvolution methods using sc/snRNAseq data as reference, such as Bisque and MuSiC, did not outperform old methods using pooled-cell reference. Given that Bisque learns prior information from the reference of sc/snRNAseq data, it is not surprising to see that Bisque showed perfect performance when using cell proportions from sc/snRNAseq data as ground truth. However, the cell proportions measured by single-cell technologies can be easily biased by the sorting strategy(*40, 41*). The proportions from single-cell data as prior reference and ground truth should be used with caution.

Cell-type expressions deconvoluted to individual samples from brain tissues were further evaluated for the first time here for eQTL mapping and differential expression analysis, which require sample-wise expression data. The development of sample-wise deconvolution satisfies these needs. Sample-wise deconvolution can estimate cell-type expression for each sample, without cell sorting and sc/snRNAseq. bMIND was the best method for estimating cell-type expressions in our evaluation, since the correlation coefficients between estimated expressions by bMIND and ground truth were higher than other methods. Moreover, our evaluation showed that the deconvoluted data by bMIND have good cell-type specificity. The deconvoluted expressions by bMIND had a high correlation with matched cell types but a low correlation with other cell types in the ground truth data (Fig. S3). The deconvoluted cell types by bMIND expressed well-known marker genes. For example, NRGN and GAD1 were highly expressed in excitatory and inhibitory neurons respectively, but poorly expressed in glial cell types. These results indicated that bMIND is the best method for generating cell-type-specific gene expression data for each sample directly from bulk tissue data.

The deconvolution performance on rare cell types is in general poor and capturing rare cell types is thus a challenge for cell deconvolution. The accuracy of deconvoluting cell proportions decreased sharply when cell proportion was less than 5%. Similarly, the accuracy of estimated gene expressions was low for rare cell types in brains, such as endothelial cells (correlation coefficient between deconvoluted expressions and ground truth = 0.33) and pericytes (correlation coefficient = 0.43). The low abundance of rare cell types may be masked by dominant cell types in the bulk tissue. Rare cell types may need to be studied using RNAseq of sorted cells, or high coverage sc/snRNAseq, or techniques that enrich for rare cell types.

The most important benefit of cell deconvolution was that, more cell-type eQTLs were identified using deconvoluted data than using single-cell data and bulk-tissue data. Cell-type eQTLs have been generated with sc/snRNAseq data(*42, 43*). However, the number of individuals profiled was typically limited(*11*). To date, a total of 7,607 eQTLs have been identified in the largest single-cell eQTL study(*38*) comprising 192 human brain samples. Besides the small sample size, the quality of single-cell data can be potentially affected by poor expression quantification, with serious dropout issues and high technical variability(*9*). Consequently, the eQTLs identified from sc/snRNAseq data may be affected. In contrast, this study deconvoluted data from 861 human brain and mapped 25,273 decon-eQTLs, far more than eQTLs identified from single-cell studies. The union for decon-eQTLs of all cell types was more than bulk-tissue eQTLs (n=9,148). A total of 24.3% ∼ 36.89% of top decon-eQTLs were not detected by bulk eQTL mapping. This indicates that many cell-type-specific eQTLs are buried in bulk-tissue data since the expression of diverse cell types is mixed. Overall, decon-eQTLs could be replicated in bulk-tissue eQTLs (averaged Pi1=0.99) and single-cell eQTLs (averaged Pi1=0.38), indicating the reliability of eQTLs identified in deconvoluted data. Sample-wise deconvolution provides a valuable opportunity to study genetic regulations in specific cell types with comparable power to bulk-tissue eQTL studies.

Decon-eQTLs explained SCZ GWAS heritability that was missed by single-cell and bulk-tissue eQTLs. Nearly six times more SCZ GWAS heritability was explained by decon-eQTLs than by single-cell eQTLs. Integrating decon-eQTLs and bulk-tissue eQTLs explained 63% of SCZ GWAS heritability, which was 14% more than heritability explained only by bulk-tissue eQTLs. These results suggested that SCZ GWAS risk may be mediated by genetic regulations in specific cell types, and such an effect can be captured by deconvoluted data.

Risk genes associated with SCZ GWAS can be revealed by decon-eQTL mapping. Identification of genetically risk genes and pathways in specific cell types is an essential application of decon-eQTLs. For example, we identified that the association between rs12466331 and CALM2 was significant in excitatory neurons but not in bulk-tissue data (Fig. S8A). CALM2, a gene encoding calmodulin, is highly expressed in excitatory neurons (Fig. S8B) and has been found downregulated in the postmortem brains of SCZ patients(*44*). Moreover, rs12466331 was colocalized with SCZ GWAS risk locus rs144040771 (Fig. S8C). These data suggest that rs12466331 may regulate the expression of CALM2 in excitatory neurons and that dysregulation of such pathway may be associated with SCZ. Thus, mapping decon-eQTLs enabled the discovery of the genetic risk of disease and helped identify their molecular mechanisms in specific cell types.

Cell-type eQTLs mapping with deconvoluted data is an advanced alternative for ieQTL mapping. ieQTLs are the results of interaction between genetic regulation and cell-type enrichment, while decon-eQTLs are based on deconvoluted data, which are the direct relationship between genotypes and cell type expression for each SNP-gene pair. Both ieQTLs and decon-eQTLs were mapped with ROSMAP data in the current study. Nearly twenty times more decon-eQTLs (n=27,339) were identified than ieQTLs (n=1,822) for the same sample size. Moreover, decon-eQTL is more robust than ieQTL. Compared to single-cell eQTLs, the replication rate of decon-eQTLs (averaged Pi1=0.38) is clearly superior than ieQTLs (averaged Pi1=0.16, Fig.S9).

This study offers a practical guideline for conducting brain cell deconvolution. Using dtangle to estimate cell proportions and bMIND to estimate cell-type expressions is recommended. Rare cell types (proportion<5%) are not recommended to be included in cell deconvolution analysis.

This study has several limitations. The results were based on the analysis of human brain data with specific parameters tested. More tests may be needed to generalize the conclusion to other tissues and situations. Unsupervised deconvolution methods have not been evaluated by this study. This evaluation only focused on the major cell types in brains, and the deconvolution performance of cell subtypes could be further explored to validate our findings.

## Conclusion

This study comprehensively evaluated the commonlu-used methods for sample-wise deconvolution of cell proportions and cell type gene expressions. The downstream analysis of eQTL mapping, GWAS heritability enrichment, and differential expression was also evaluated. Our analysis is a crucial methodological foundation for other studies where deconvolution can be used. A practical guideline is offered for a broad community interested in cell-type-specific studies of brain functions and disorders when only bulk-tissue transcriptome is available.

## Materials and Methods

### Data processing

#### Bulk-tissue RNAseq data

Three RNAseq data from brain tissues and brain organoids were used (Table 1). TMM normalization(*45*) was applied to the raw counts data and log-transformed counts per million reads mapped (CPM) were used. Gene with log2CPM>0.1 in at least 25% of samples were retained. Connectivity between samples was calculated by weighted correlation network analysis (WGCNA)(*46*) and z-score was normalized. Samples with z-score connectivity < (-3) were labeled as outliers and were removed from downstream analysis. Data were then quantile normalized with the preprocessCore(*47*) package. The batch effect was corrected with combat in the sva package(*48*).

#### Sc/snRNAseq data

The processed count matrix and metadata were used. The ROSMAP snRNAseq data were downloaded from https://www.synapse.org/#!Synapse:syn18681734. For CMC snRNAseq data, the processing pipeline can be found in the Capstone paper (syn48958066). scRNAseq data of brain organoids can be found in Jourdon et al(*35*).

#### IHC data

IHC data were downloaded from https://github.com/ellispatrick/CortexCellDeconv. Cell proportions were normalized according to the sum-to-1 constraint.

### Construction of references and pseudo-bulks

Two types of references were used. For DSA, dtangle, OLS, and CIBERSORT, pooled-cell reference is required. To build a pooled-cell reference, the count matrix was averaged by cell types. The averaged counts matrix was normalized into CPM and was log2-transformed. For MuSiC and Bisque, a single-cell reference was used, which was the gene-by-cell count matrix. To build pseudo-bulks, the count matrix was summed by cell types and by individuals.

### Marker gene identification

Marker genes were identified at the cell level and pseudo-bulk level. One versus second high strategy was used. For each gene, the expression difference between the cell type with the highest expression and the cell type with the second highest expression was calculated. At the cell level, marker genes were identified with Seurat(*49*). Genes having a proportion of zero expression>15% in the target cell type were removed. The Wilcoxon signed-rank test was used to test the expression difference. Genes with log2FC>1 and FDR corrected p value<0.05 were defined as marker genes at the cell level. At the pseudo-bulk level, marker genes were tested in DESeq2(*50*). The likelihood ratio test was used to test the expression difference between the two cell groups. Marker genes with log2FC>2 and FDR-corrected p-value <0.05 were defined as marker genes at the pseudo-bulk level.

### Estimation of cell proportions

Three inputs were required for all deconvolution methods: bulk tissue data, reference, and marker genes. Batch-corrected data was used as input for bulk tissue data. The intersected genes between marker genes at the cell level and pseudo-bulk level were used as input of marker genes. For DSA, dtangle, OLS, and CIBERSORT, pooled-cell reference was used. For MuSiC and Bisque, single-cell reference was used. The genes that have no expression variation were removed from the reference.

### Evaluation of cell proportions

Two ground truths were used to evaluate estimated cell proportions: cell proportions from IHC data and cell proportions from sc/snRNAseq data. For one sample, cell proportions from sc/snRNAseq were calculated by dividing the number of cells of one specific cell type by the total number of cells. RMSE was used as an evaluation metric. The formula of RMSE is:

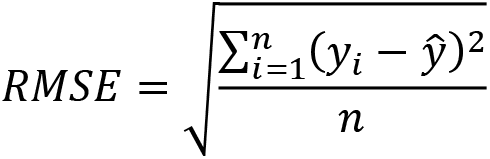

*y_i_* is the estimated cell proportion and *ŷ* is the ground truth. n is the number of cell types in the sample-level evaluation, and n is the number of samples in the cell-type-level evaluation.

### Estimation of cell-type expression

Batch-corrected data was used as input for bulk tissue data. The proportion from DSA, dtangle, OLS, CIBERSORT, MuSiC, and Bisque was used independently. Pooled-cell reference was used as prior for bMIND and swCAM. For TCA, only bulk-tissue data and cell proportions were used to estimate cell type expressions for each sample. Data were transformed into a log scale for bMIND and TCA and a linear scale for swCAM.

### Evaluation of cell type expression

To construct ground truth for evaluating estimated expressions, averaged counts by cell types in each sample were calculated. Then the averaged cell type expressions were normalized into CPM and were log2-transformed. Sample-to-sample spearman correlation was tested between estimated expression and ground truth for each cell type.

### Genotyping quality control

The ROSMAP whole genome sequencing (WGS) dataset was downloaded from https://www.synapse.org/#!Synapse:syn11724057. The data is already imputed. Only individuals with both genotype and deconvolution results were retained for the eQTL analysis. SNPs with minor allele frequency (MAF) <5% or deviating from Hardy– Weinberg equilibrium (P < 1 × 10-6) were excluded. After quality control, we obtained high-quality genotypes for ∼4.9 million SNPs (MAF > 5%) in 861 individuals.

### eQTL mapping

#### decon-eQTLs

To identify decon-eQTLs, we tested the associations between genotypes and deconvoluted expressions. We mapped cis-eQTLs within a 1-Mb window of the TSS of each gene using QTLtools(*51*). For each gene, QTLtools performs permutations of the expression data and records the best p-value for each SNP in the cis window after each permutation. We used estimated cell-type expression by bMIND as phenotype data. Phenotype data of eight cell types were tested independently. Quantile normalization was used for normalizing expression matrixes before eQTL mapping. PEER was used to identify hidden covariates in the data(*52*). 8-35 PEER factors were included as covariates in eQTL mapping.

#### Bulk-tissue eQTLs

To map bulk-tissue eQTLs, the same eQTL mapping procedure was performed on the bulk-tissue expression data. 33 PEER factors were included as covariates in eQTL mapping.

### Replication of decon-eQTLs in BrainGVEX data

To replicate decon-eQTLs, we deconvoluted RNAseq data from BrainGVEX(*36*) and mapped eQTLs with the deconvoluted data. 430 brain samples with both genotypes and RNAseq data were used. dtangle was used to estimate cell proportions, with the marker genes and the reference from ROSMAP sn/RNAseq data. Then, bulk-tissue data were deconvoluted into cell-type expressions for eight major cell types with bMIND. The same eQTL mapping process was performed on the deconvoluted data to identify decon-eQTLs in BrainGVEX data.

The proportion of true associations (π1) in the qvalue package(*37*) was used to measure the replicate rate of significant decon-eQTLs in ROSMAP data in decon-eQTLs in BrainGVEX data. With the distribution of corresponding p values for the overlapped eSNP-eGene pairs in two datasets, we calculated π0, i.e., the proportion of true null associations based on distribution. Then, π1 = 1 - π0 estimated the lowest bound for true-positive associations.

### Replication of decon-eQTLs in single-cell eQTLs

To measure the replication rate of decon-eQTLs in the sc/snRNAseq dataset, we downloaded cell-type eQTLs identified from the snRNAseq data of 192 individuals(*38*). With the single-cell eQTLs as a reference, π1 statistics were calculated for eight cell types independently.

### SCZ heritability enrichment

Stratified linkage disequilibrium score regression(*39*) (S-LDSC) was used to calculate SCZ GWAS heritability enrichment in decon-eQTLs. GWAS summary statistics from three published SCZ studies were downloaded(*53-55*). Conditional analysis was performed on decon-eQTLs to select the top SNP for each gene (r2 > 0.2 in 1000 Genomes European individuals(*56*)). Then script ldsc.py with the “--l2” parameter was used to generate the gene-set-specific annotation and LD score files. Then ldsc.py with the “--h^2^-cts” parameter was used to generate stratified heritability by decon-eQTLs of eight cell types.

### Co-localization

For each gene in decon-eQTLs, the co-localization between eSNP and SCZ GWAS signals(*57*) was tested. The ‘coloc.abf’ function in the Coloc(*58*) package (version 5.1.0) was used for testing. The threshold for significance is SNP.PP.H4>0.95.

### Differential expression analysis

Differential expression analysis was performed on deconvoluted data and bulk-tissue data to identify genes associated with AD, SCZ, and brain development. For AD and SCZ, differential expression analysis was conducted in each cell type with the Wilcoxon rank-sum test. For brain development, the linear regression model was used to identify genes showing significant expression changes. The p values were corrected by FDR. Genes with FDR q value <0.05 were identified as phenotype-associated genes (PAGs).

To compare deconvoluted PAGs and PAGs from sc/snRNAseq data, PAGs for AD(*23*) and SCZ(*59*) in cell types were downloaded. For brain development, PAGs were identified in pseudo-bulk data. The linear regression model was used to identify PAGs for each cell type independently. The p values were corrected by FDR.

## Data availability

The source data described in this manuscript are available via the PsychENCODE Knowledge Portal (https://psychencode.synapse.org/). The PsychENCODE Knowledge Portal is a platform for accessing data, analyses, and tools generated through grants funded by the National Institute of Mental Health (NIMH) PsychENCODE Consortium. Data is available for general research use according to the following requirements for data access and data attribution: (https://psychencode.synapse.org/DataAccess). For access to content described in this manuscript see: https://www.synapse.org/#!Synapse:syn51072187/datasets/. The eQTL and PAG results can be accessed at https://www.synapse.org/#!Synapse:syn50908925.

## Acknowledgment

We thank Richard Kopp at SUNY Upstate Medical University for his help in polishing words. We thank all the participants involved in the ROSMAP and PsychENCODE study for making the data available. This work was supported by NIH grants U01MH122591, U01MH116489, R01MH110920, U01MH103340, R01MH126459 and R01MH109648; the Simons Foundation 632742; the National Natural Science Foundation of China 82022024 and 31970572; the science and technology innovation Program of Hunan Province 2021RC4018 and 2021RC5027. Data were generated as part of the PsychENCODE Consortium, supported by: U01DA048279, U01MH103339, U01MH103340, U01MH103346, U01MH103365, U01MH103392, U01MH116438, U01MH116441, U01MH116442, U01MH116488, U01MH116489, U01MH116492, U01MH122590, U01MH122591, U01MH122592, U01MH122849, U01MH122678, U01MH122681, U01MH116487, U01MH122509, R01MH094714, R01MH105472, R01MH105898, R01MH109677, R01MH109715, R01MH110905, R01MH110920, R01MH110921, R01MH110926, R01MH110927, R01MH110928, R01MH111721, R01MH117291, R01MH117292, R01MH117293, R21MH102791, R21MH103877, R21MH105853, R21MH105881, R21MH109956, R56MH114899, R56MH114901, R56MH114911, R01MH125516, R01MH126459, R01MH129301, R01MH126393, R01MH121521, R01MH116529, R01MH129817, R01MH117406, and P50MH106934 awarded to: Alexej Abyzov, Nadav Ahituv, Schahram Akbarian, Kristin Brennand, Andrew Chess, Gregory Cooper, Gregory Crawford, Stella Dracheva, Peggy Farnham, Michael Gandal, Mark Gerstein, Daniel Geschwind, Fernando Goes, Joachim F. Hallmayer, Vahram Haroutunian, Thomas M. Hyde, Andrew Jaffe, Peng Jin, Manolis Kellis, Joel Kleinman, James A. Knowles, Arnold Kriegstein, Chunyu Liu, Christopher E. Mason, Keri Martinowich, Eran Mukamel, Richard Myers, Charles Nemeroff, Mette Peters, Dalila Pinto, Katherine Pollard, Kerry Ressler, Panos Roussos, Stephan Sanders, Nenad Sestan, Pamela Sklar, Michael P. Snyder, Matthew State, Jason Stein, Patrick Sullivan, Alexander E. Urban, Flora Vaccarino, Stephen Warren, Daniel Weinberger, Sherman Weissman, Zhiping Weng, Kevin White, A. Jeremy Willsey, Hyejung Won, and Peter Zandi. The authors declare that they have no competing interests.

## Supplemental Figures

**Fig. S1.**
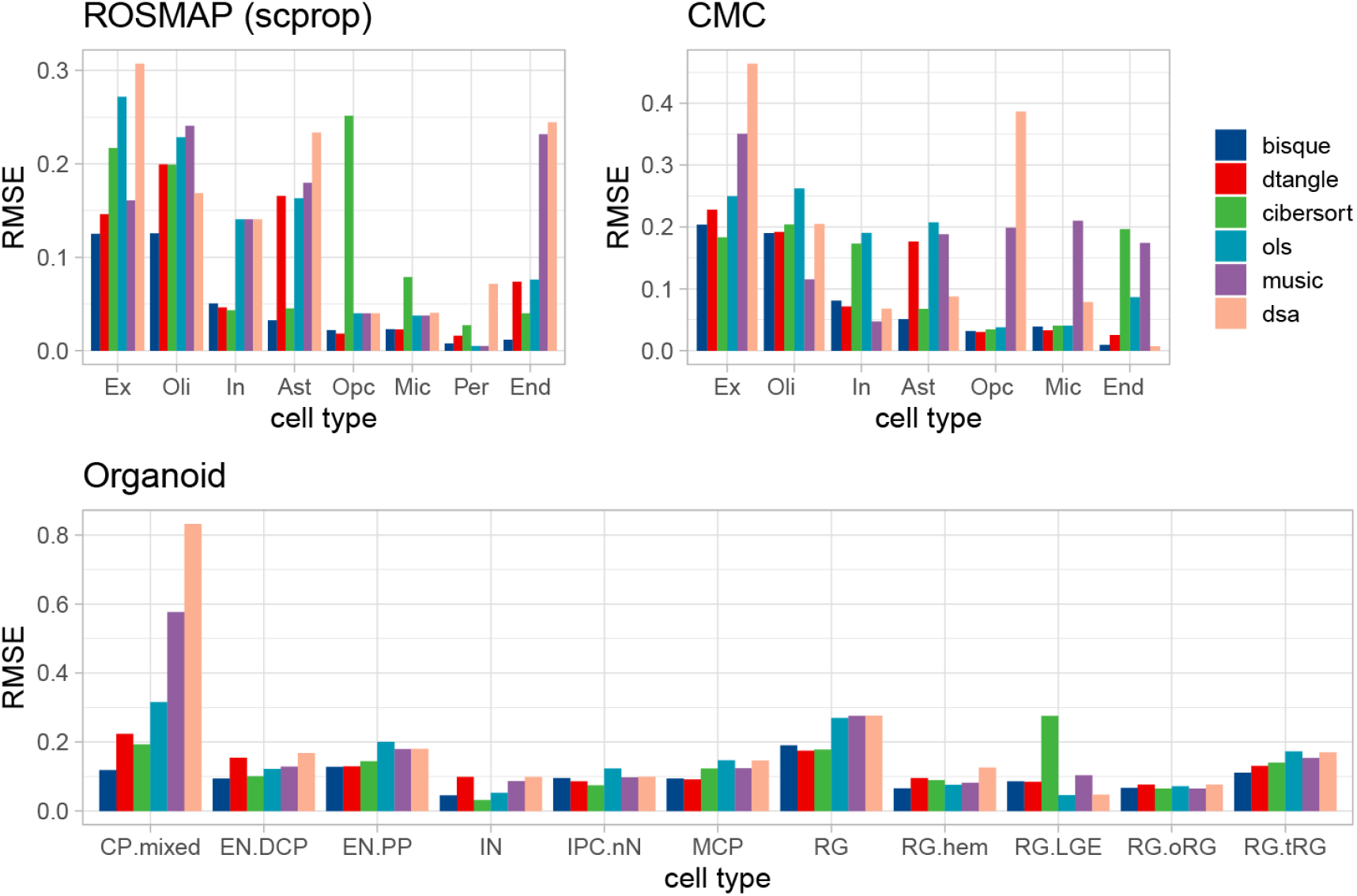
Cell-type-level RMSE values between estimated cell proportions and ground truth. This is the full version of Fig. 2B in terms of RMSE values.

**Fig. S2.**
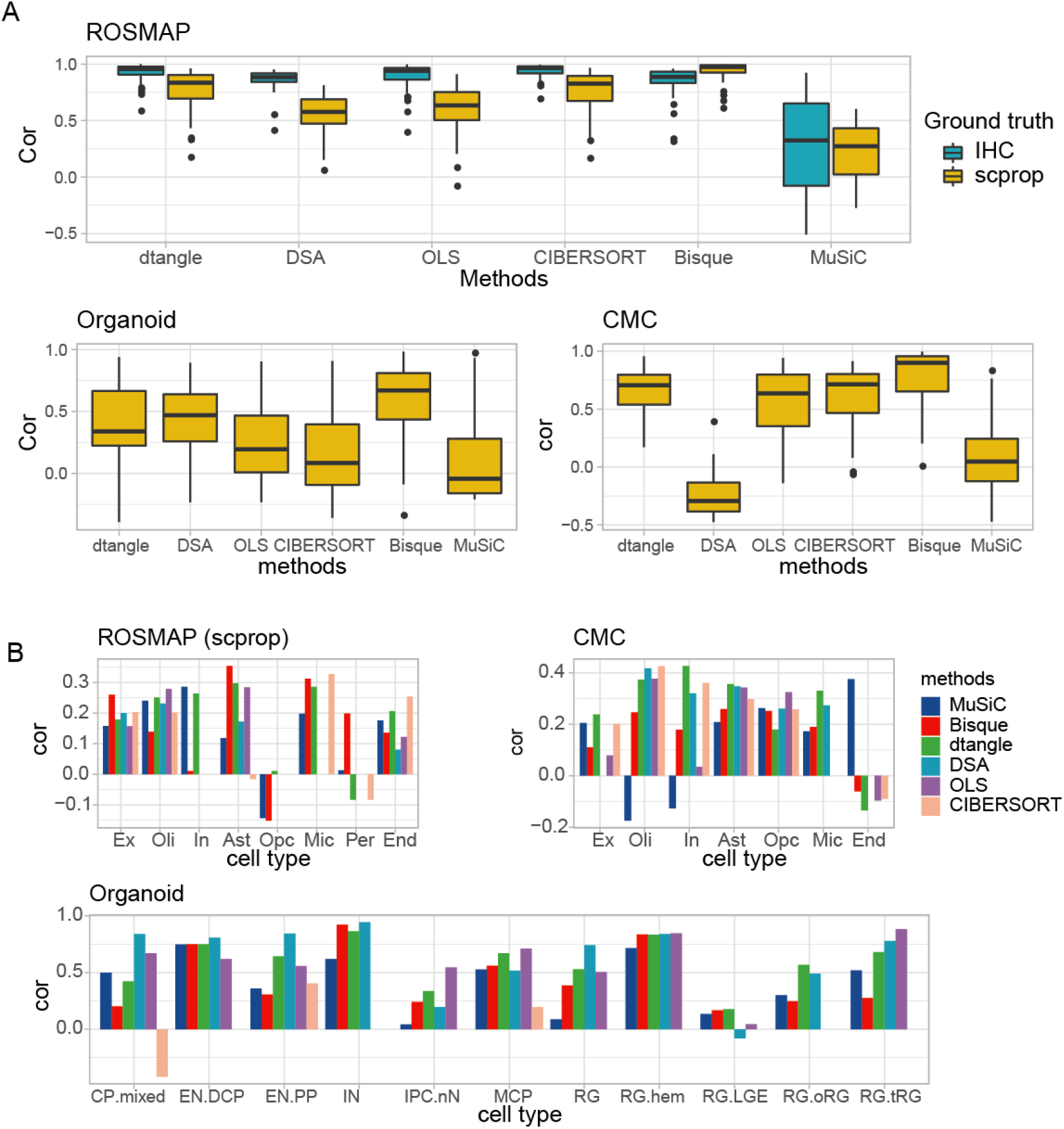
Assessment of cell proportions estimated by deconvolution methods based on Spearman correlation. **(A).** The sample-level correlation coefficient between estimated cell proportions and ground truth. IHC: immunohistochemistry; scprop: cell proportions calculated from sc/snRNAseq data, scprop = the number of cells of specific cell type/number of total cells. **(B).** The cell-type-level correlation coefficient between estimated cell proportions and ground truth. Cell types were ordered by cell proportions in a decreasing way. Ex: excitatory neurons, In: inhibitory neurons, Ast: astrocytes, Opc: oligodendrocyte precursor cells, Mic: microglia, Per: pericytes, End: endothelial cells; RG: radial glia, EN.PP: early born excitatory neurons of the pre-plate/subplate, CP.mixed: cortical plate mixed neurons, MCP: medial cortical plate, EN-DCP: dorsal cortical plate excitatory neurons, IPC-nN: intermediate progenitor cell or newborn neuron, RG.tRG: truncated radial glia, RG.oRG: outer radial glia, RG.hem: radial glia in cortical hem, IN: inhibitory neurons, RG-LGE: progenitors corresponding to a putative ventrolateral ganglionic eminence fate.

**Fig. S3.**
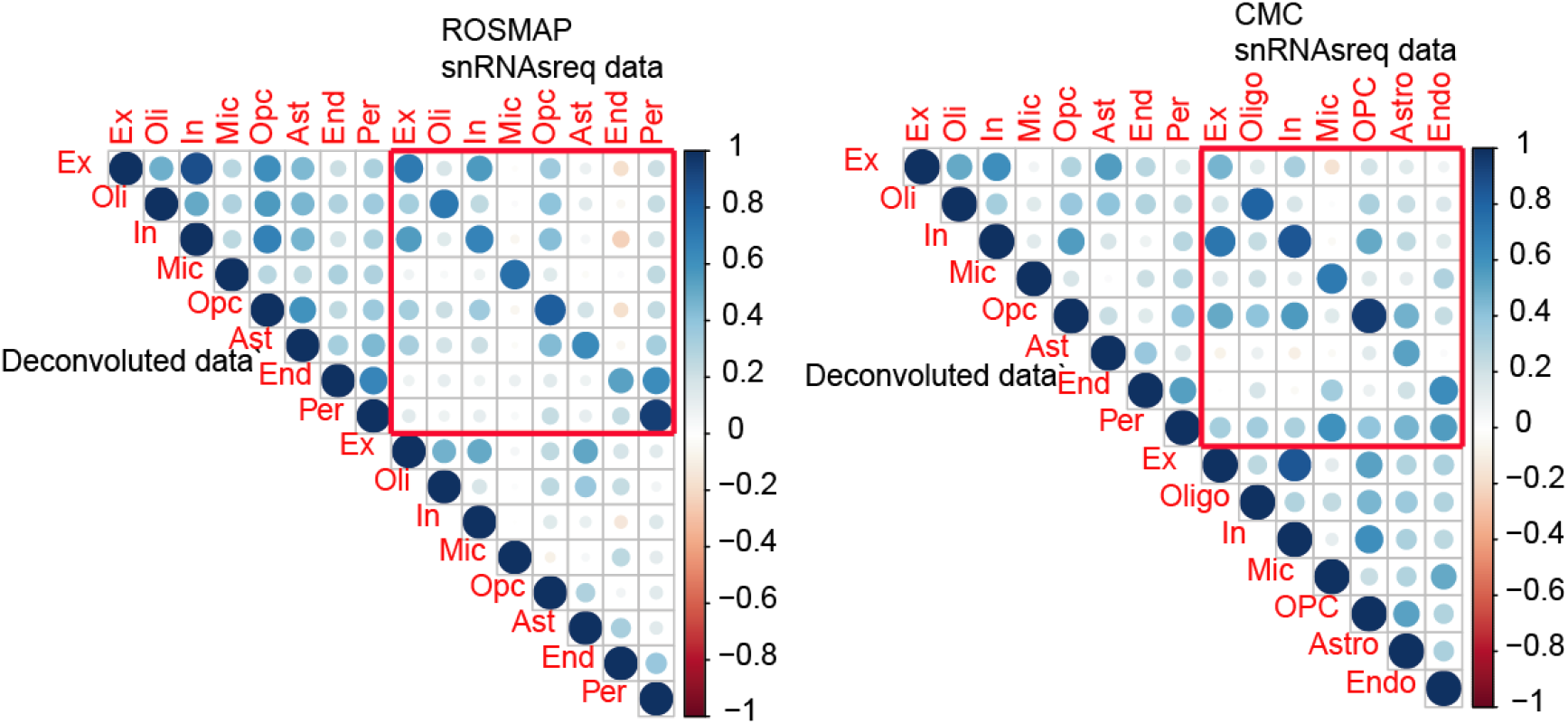
Correlations between deconvoluted expression (ROSMAP) and snRNAseq data from ROSMAP and CMC. For each cell type, averaged expressions across all samples were used.

**Fig. S4.**
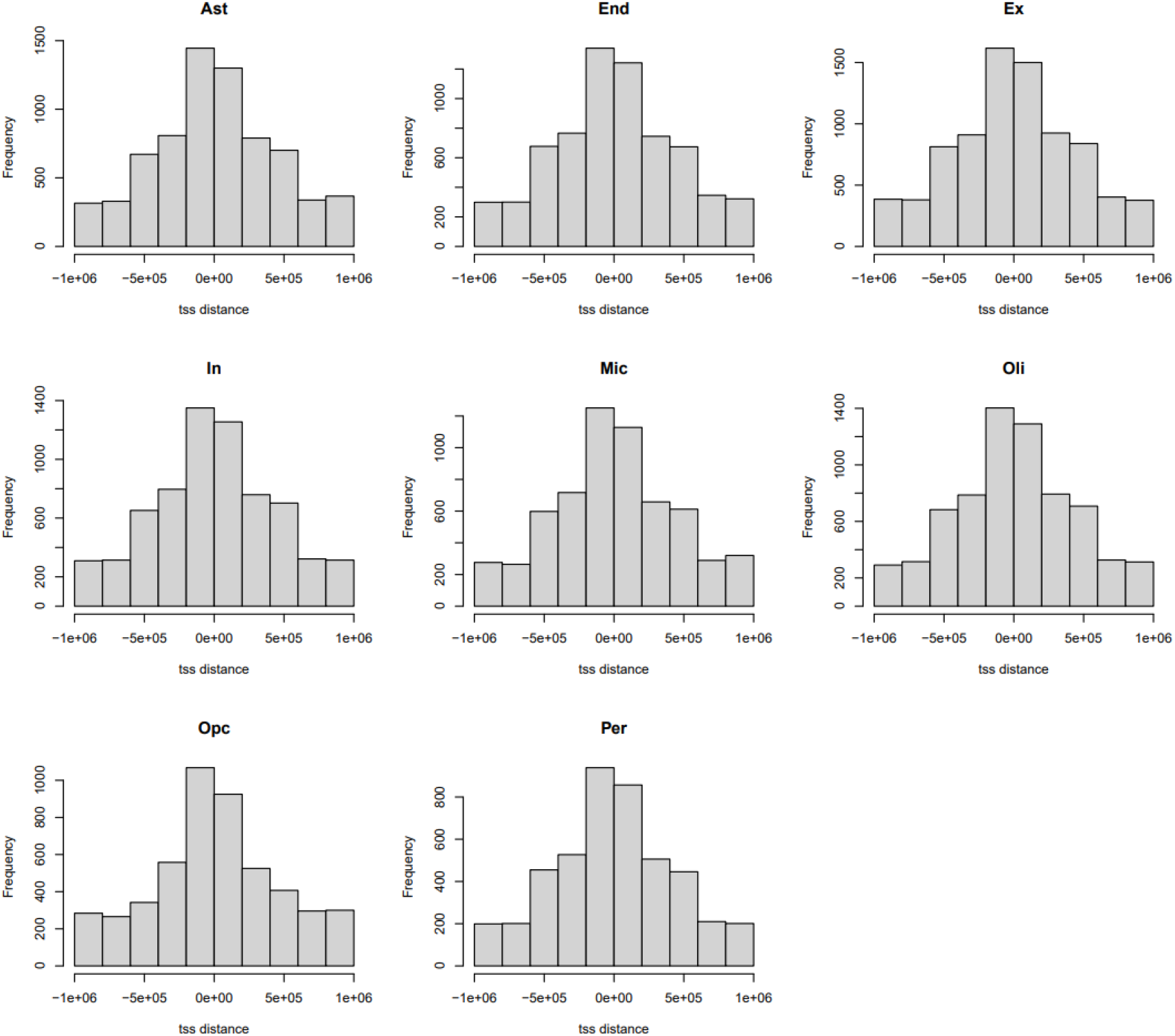
Distance between transcription start sites (TSS) and eQTL SNPs (eSNPs).

**Fig. S5.**
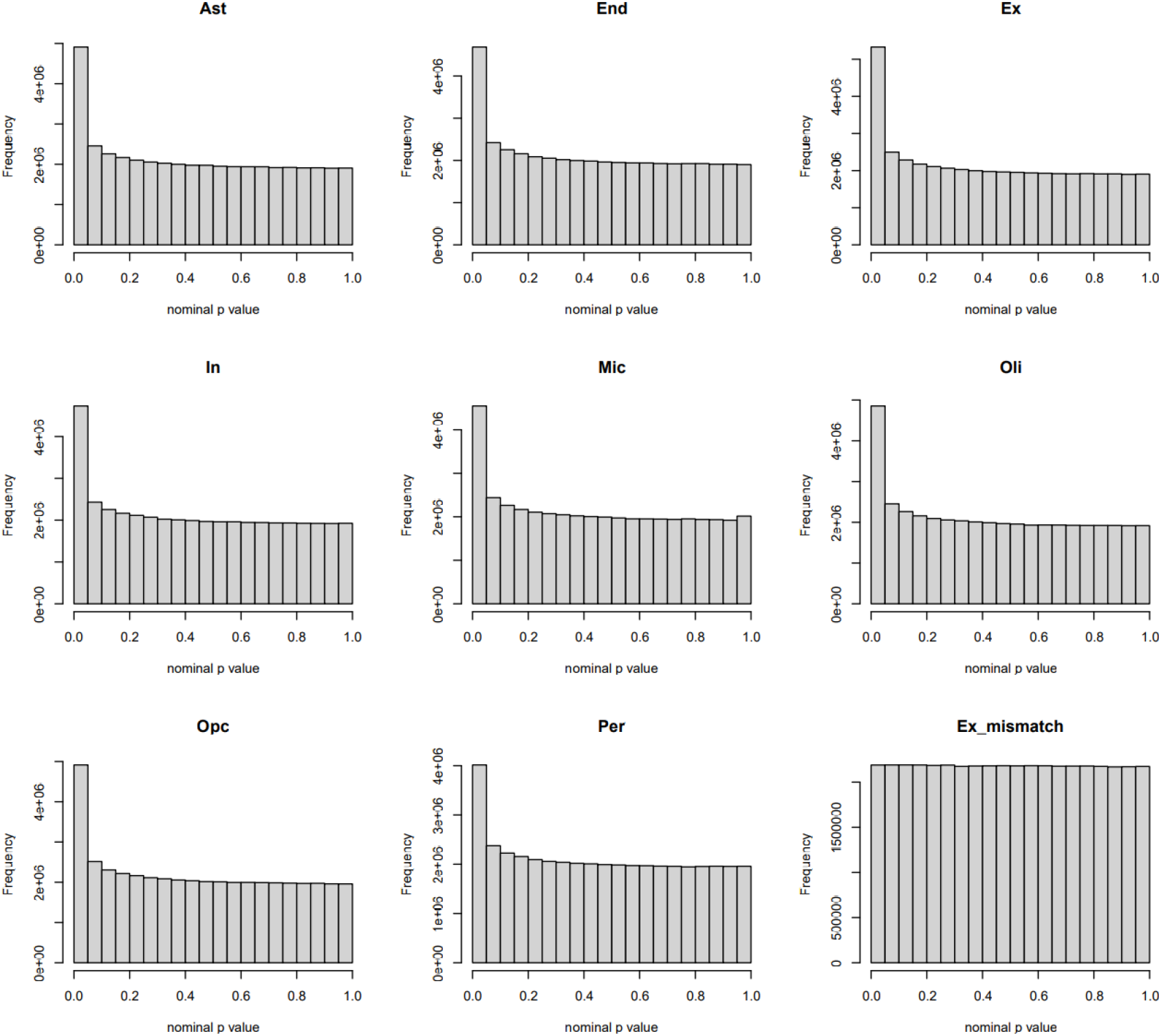
Distribution of decon-eQTL p values. Ex_mismatch represents eQTL mapping results based on sample-shuffled data of deconvoluted excitatory neurons.

**Fig. S6.**
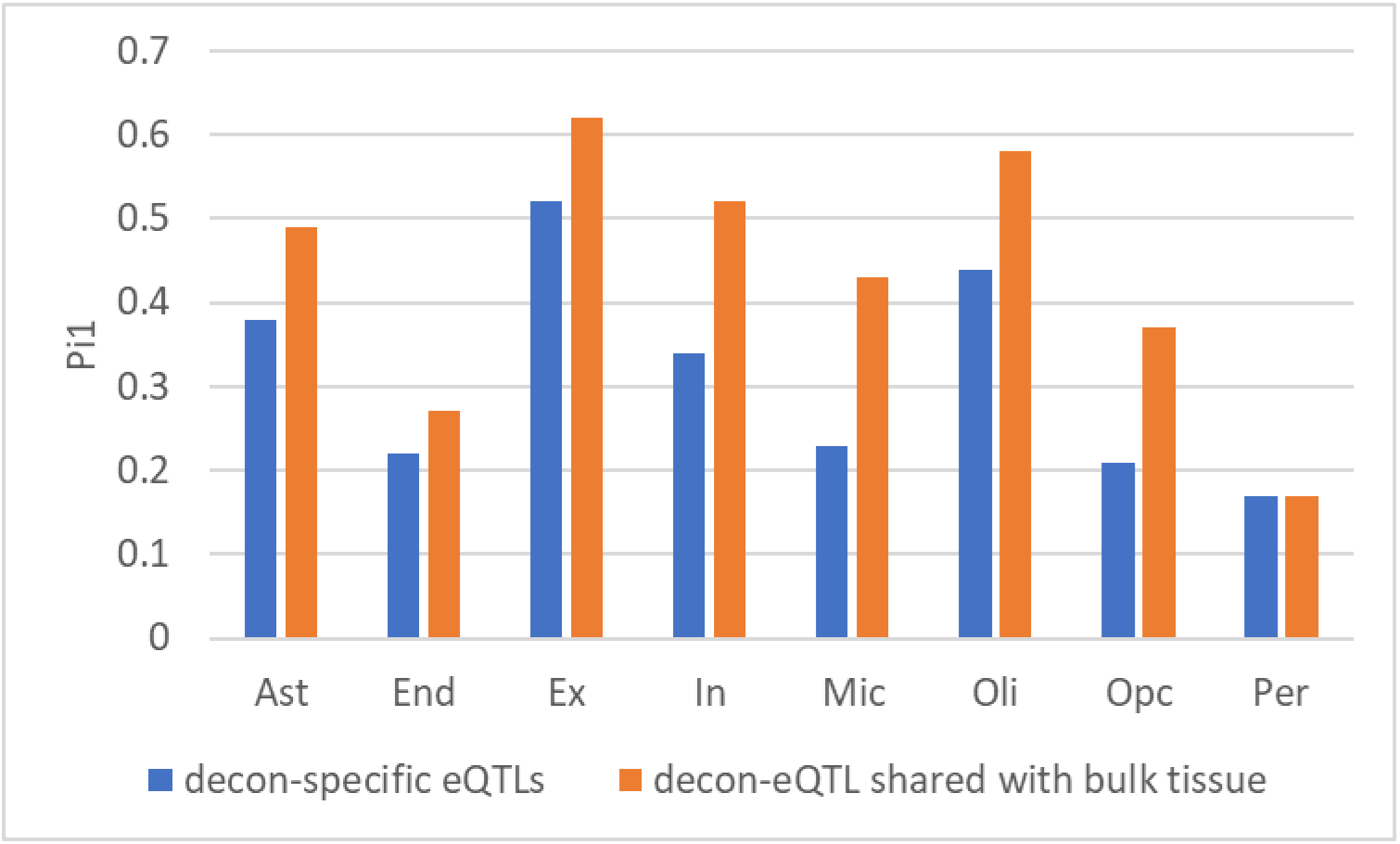
Replication of decon-eQTLs in single-cell eQTLs. Wilcoxon signed-rank test was used to test the difference in Pi1 of two eQTL classes. P=0.15.

**Fig. S7.**
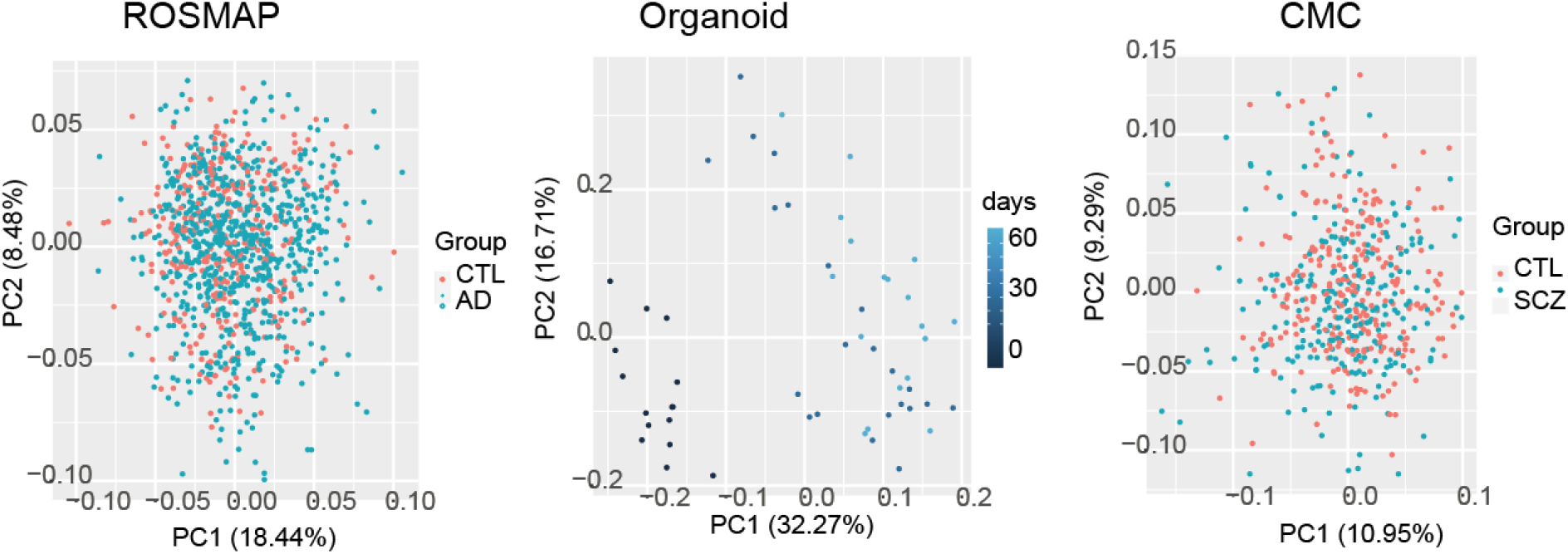
PCA plot of samples in bulk-tissue datasets. Batch-corrected were used.

**Fig. S8.**
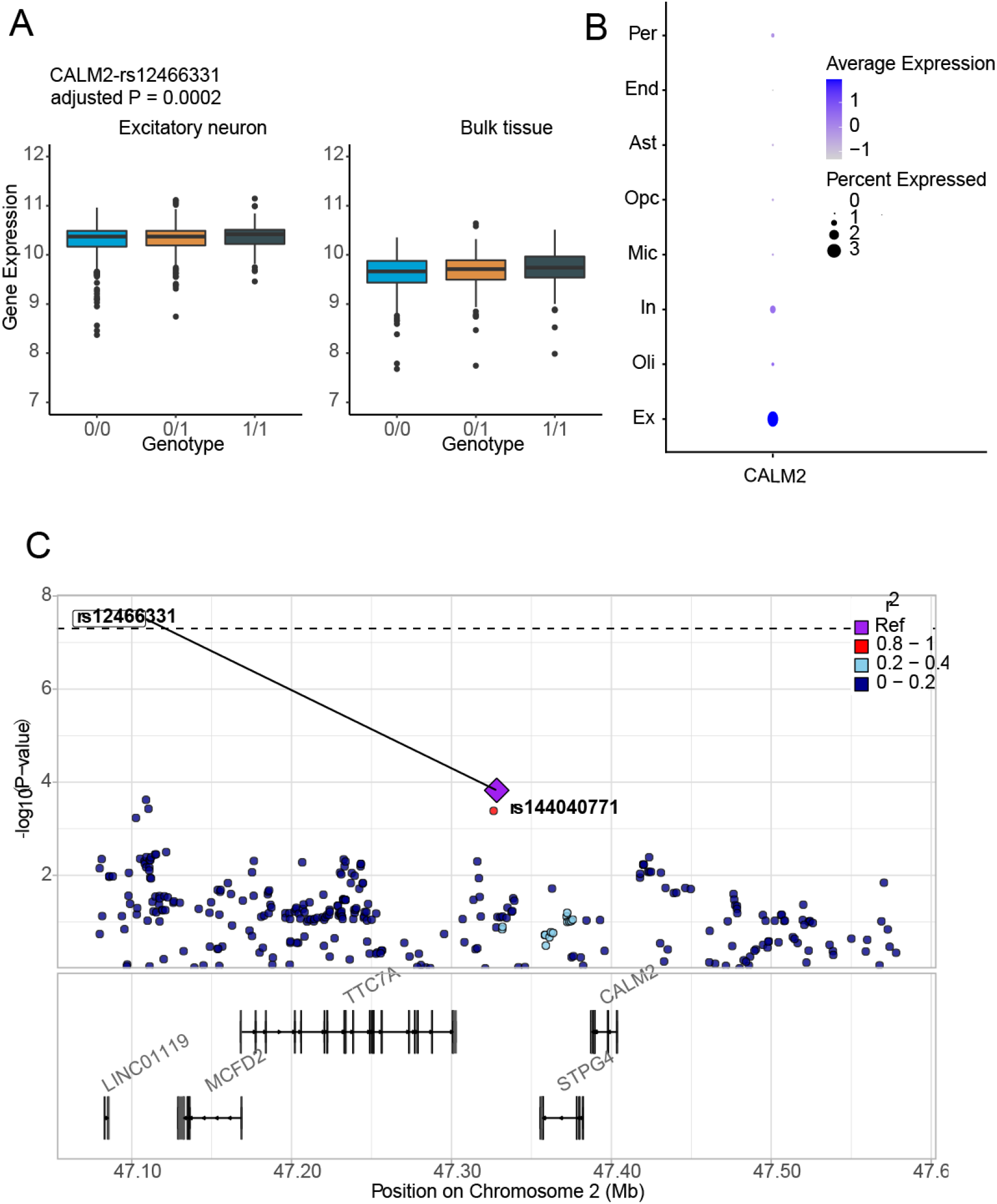
An example of cell-type specific eQTLs in excitatory neurons. **(A)** Expression of CALM2 in individuals with different genotypes. **(B)** Expression of CALM2 in ROSMAP snRNAseq data. **(C)** Colocalization of eSNPs on CALM2 and SCZ GWAS risk locus.

**Fig. S9.**
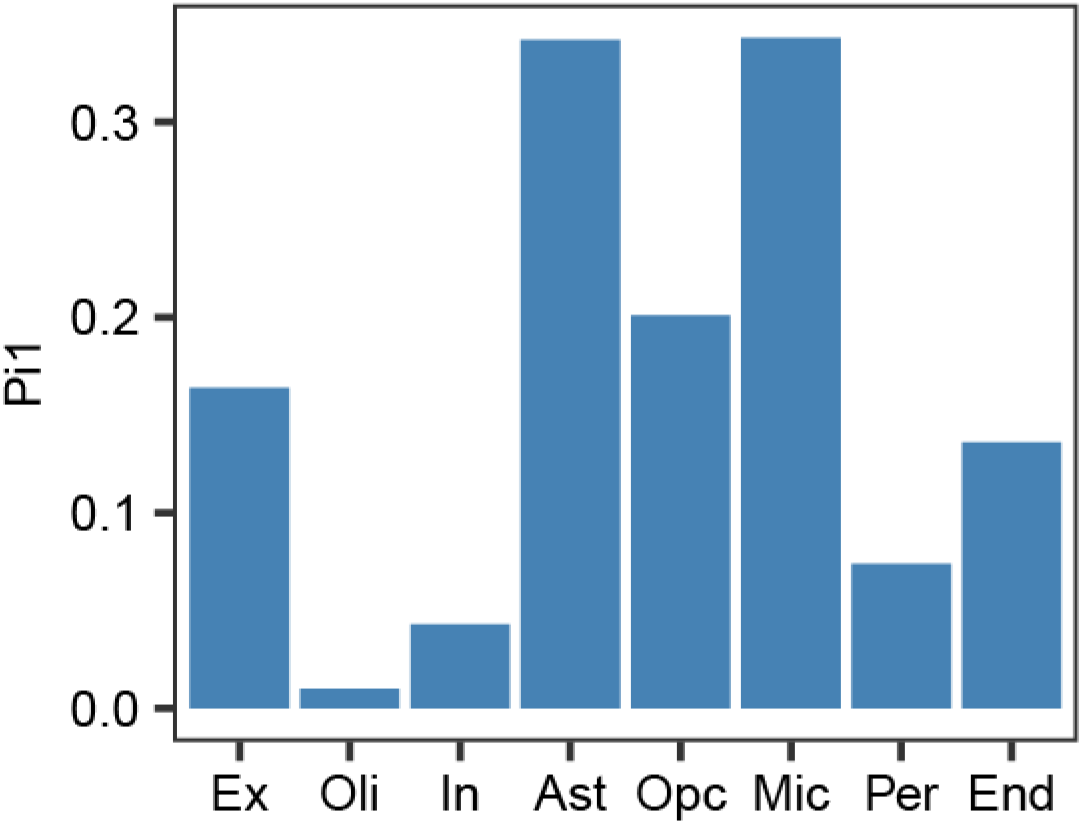
Replication of ieQTLs in single-cell eQTLs.

## Notes

### Competing Interest Statement

The authors have declared no competing interest.

